# Structure of SARS-CoV-2 ORF8, a rapidly evolving coronavirus protein implicated in immune evasion

**DOI:** 10.1101/2020.08.27.270637

**Authors:** Thomas G. Flower, Cosmo Z. Buffalo, Richard M. Hooy, Marc Allaire, Xuefeng Ren, James H. Hurley

**Affiliations:** Department of Molecular and Cell Biology and California Institute for Quantitative Biosciences, University of California, Berkeley, Berkeley, CA 94720; Molecular Biophysics & Integrated Bioimaging Division, Lawrence Berkeley National Laboratory, Berkeley, CA 94720

## Abstract

The molecular basis for the severity and rapid spread of the COVID-19 disease caused by SARS-CoV-2 is largely unknown. ORF8 is a rapidly evolving accessory protein that has been proposed to interfere with immune responses. The crystal structure of SARS-CoV-2 ORF8 was determined at 2.04 Å resolution by x-ray crystallography. The structure reveals a ~60 residue core similar to SARS-CoV ORF7a with the addition of two dimerization interfaces unique to SARS-CoV-2 ORF8. A covalent disulfide-linked dimer is formed through an N-terminal sequence specific to SARS-CoV-2, while a separate non-covalent interface is formed by another SARS-CoV-2-specific sequence, _73_YIDI_76_. Together the presence of these interfaces shows how SARS-CoV-2 ORF8 can form unique large-scale assemblies not possible for SARS-CoV, potentially mediating unique immune suppression and evasion activities.

---

The severity of the current COVID-19 pandemic caused by SARS-CoV-2 relative to past outbreaks of MERS, SARS, and other betacoronaviruses, in humans begs the question as to its molecular basis. The accessory protein ORF8 is one of the most rapidly evolving betacoronavirus proteins (*1–7*). While ORF8 expression is not strictly essential for SARS-CoV and SARS-CoV-2 replication, a 29 nucleotide deletion (Δ29) that occurred early in human to human transmission of SARS-CoV, splitting ORF8 into ORF8a and 8b, is correlated with milder disease (*8*). A 382 nucleotide deletion (Δ382) in SARS-CoV-2 (*9, 10*) was also found to correlate with milder disease and a lower incidence of hypoxia (*11*).

SARS-CoV-2 ORF8 is a 121 amino acid protein consisting of an N-terminal signal sequence followed by a predicted Ig-like fold (*12*). With <20% sequence identity to SARS-CoV ORF8, SARS-CoV-2 ORF8 is remarkably divergent. ORF8 proteins from both viruses possess a signal sequence for ER import. Within the lumen of the ER, SARS-CoV-2 ORF8 interacts with a variety of host proteins, including many factors involved in ERAD (*13*). Presumably, ORF8 is secreted, rather than retained in the ER, since ORF8 antibodies are one of the principal markers of SARS-CoV-2 infections (*14*). Several functions have been proposed for SARS-CoV-2 ORF8. ORF8 disrupts IFN-I signaling when exogenously overexpressed in cells (*15*). It has been shown that ORF8 of SARS-CoV-2, but not ORF8 or ORF8a/b of SARS-CoV, downregulates MHC-I in cells (*16*).

These observations suggest the relationship between ORF8 structure, function, and sequence variation may be pivotal for understanding the emergence of SARS-CoV-2 as a deadly human pathogen. Yet not only is there no three-dimensional structure of any ORF8 protein from any coronavirus, there are no homologs of known structure with sequence identity sufficient for a reliable alignment. SARS and SARS-CoV-2 ORF7a are the most closely related templates of known structure(*17*), yet its core is approximately half the size of ORF8 and its primary sequence identity is negligible. Therefore, we determined the crystal structure of SARS-CoV-2 ORF8. The structure confirms the expected Ig-like fold and overall similarity of the core fold to SARS-CoV ORF7a. The structure reveals two novel dimer interfaces for SARS-CoV-2 ORF8 unique relative to all but its most recent ancestors in bats. Together, our results set the foundation for elucidating essential aspects of ORF8 biology to be leveraged for the development of novel therapeutics.

We generated SARS-CoV-2 ORF8 protein by expression in *E. coli* and oxidative refolding. The structure of SARS-CoV-2 ORF8 was determined by X-ray crystallography at a resolution of 2.04 Å (Fig. 1A, B). Side-chain density was visible throughout most of the density map and an atomic model was built *ab initio* into the density (Fig. 1C). ORF8 crystallized as a covalent dimer with 3 sets of intramolecular disulfide bonds per monomer and a single intermolecular disulfide bond formed by Cys20 of each monomer (Fig. 1B). The core of each ORF8 monomer consists of a two antiparallel β-sheets (Fig. 1D, 1E). The smaller sheet consists of β2 and β5, while the larger is formed from the remaining five β-strands (Fig. 1D, 1E).

**Fig 1.**
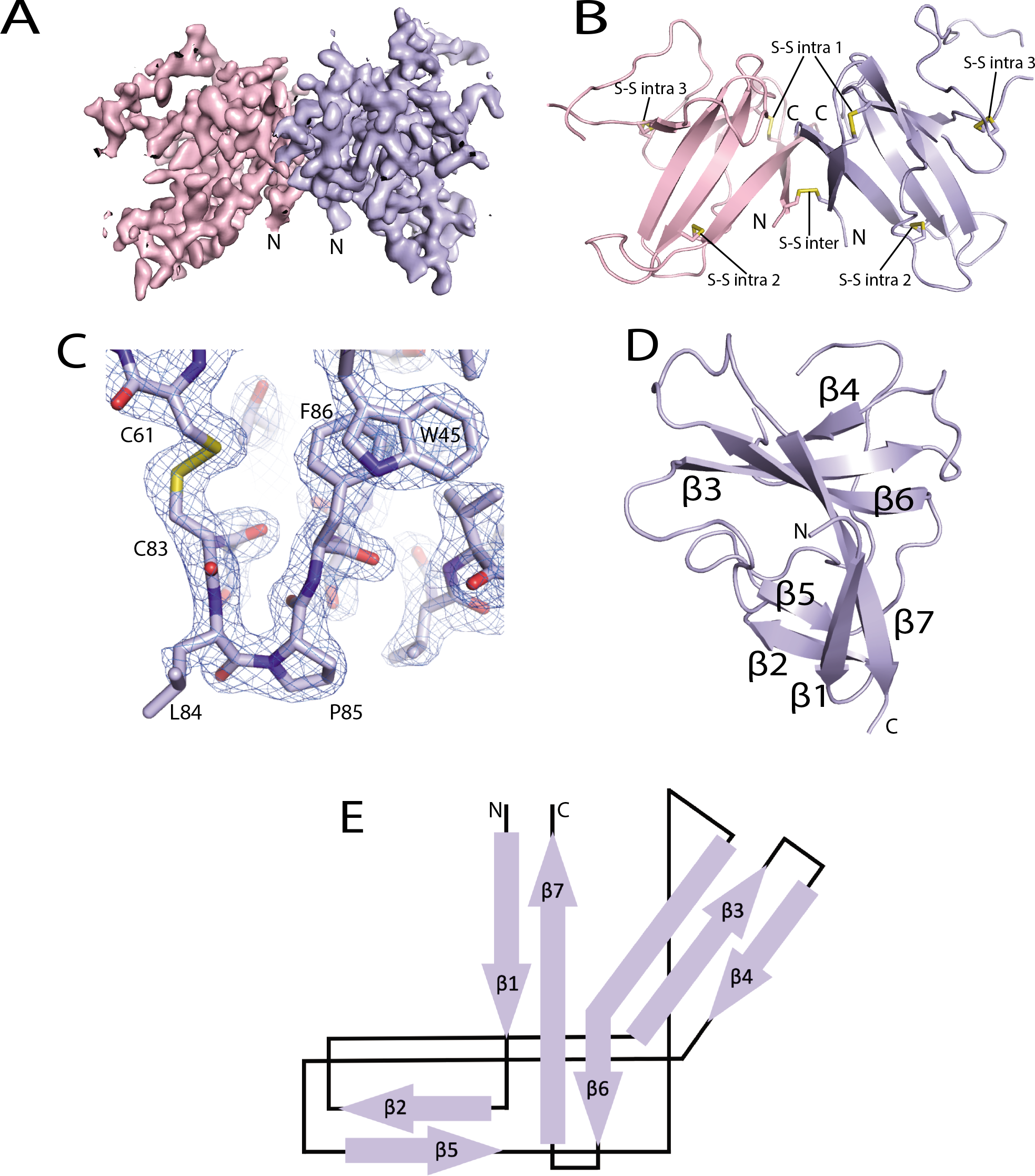
Crystal Structure of SARS-CoV-2 ORF8. (A) Electron density map of SARS-CoV-2 ORF8 crystallographic dimer determined to 2.04 Å (chain A: light blue, Chain B: light pink). (B) Cartoon representation of the SARS-CoV-2 ORF8 crystallographic dimer. Disulfide bonds are modeled showing both inter- and intramolecular bond pairs. (C) Representative density of the Cys-83-Leu84-Pro85 turn motif. The map is contoured at 2 σ and represented as a blue mesh (D) Cartoon representation of the SARS-CoV-2 ORF8 monomer. β-strands are labeled β1-β7. (E) Topographic representation of showing antiparallel β-sheets formed by β1-β7.

ORF8 has a 14% sequence identity with the SARS-CoV ORF7a protein. The Ig-like fold of ORF7a (PDB:1XAK) aligns with the ORF8 monomer with a DALI Z-score=4.6 and an RMSD=2.4 (Fig. 2B). Based on the structural alignment, ORF8 and ORF7a share two sets of structural disulfide linkages that are central to the Ig-like fold (Fig 2A, C). Punctuated between what would be β3 and β5 of ORF7a is a ORF8-specific region of ~35AA from residues 46-83 that are structurally distinct from ORF7a and other Ig-like folds (Fig. 2A, B, D). These residues are responsible for a third, ORF8-specific, disulfide (Fig 2A, C).

**Fig 2.**
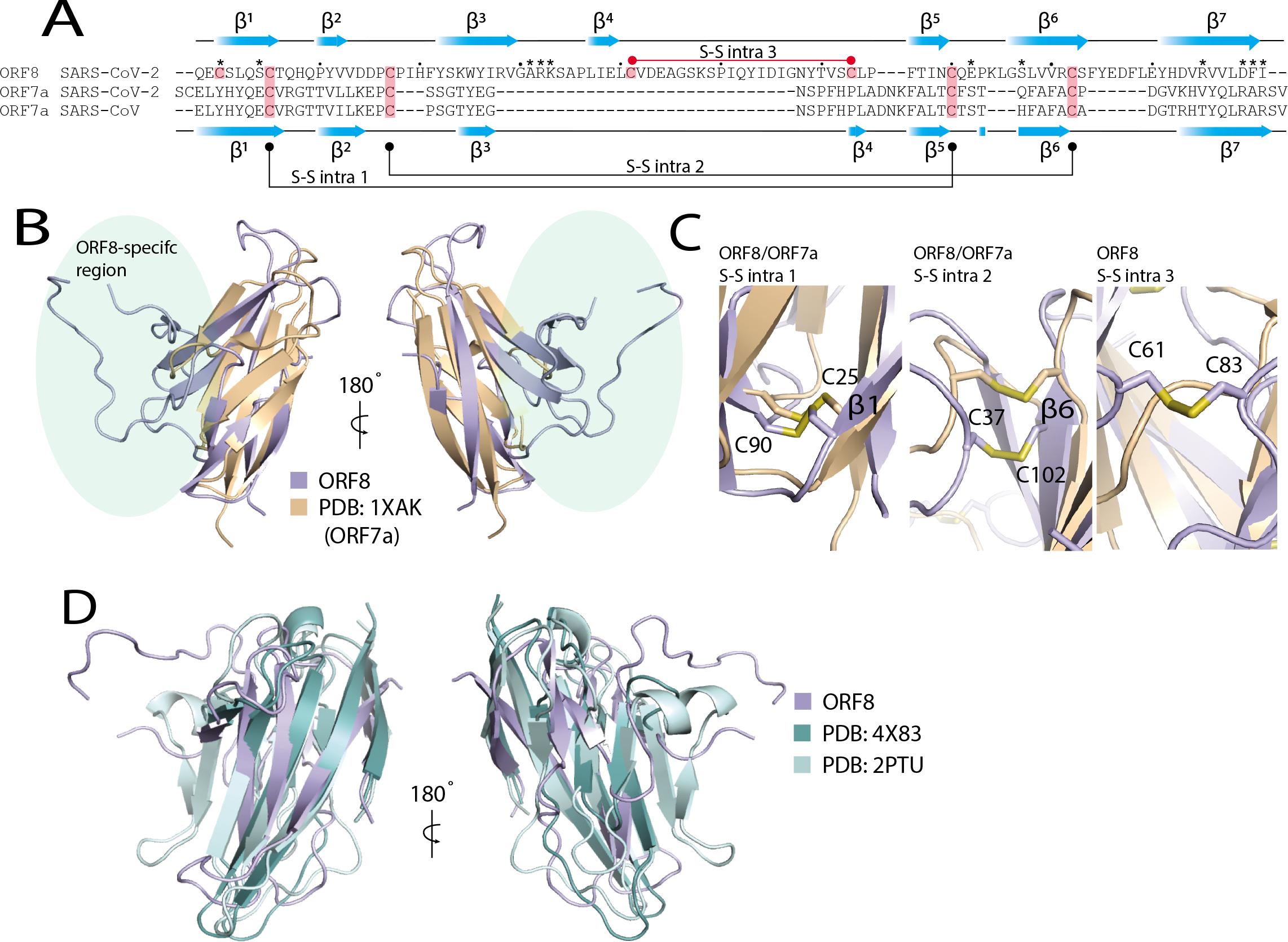
SARS-CoV-2 ORF8 adopts an Ig-like fold. (A) Structure-guided sequence alignment of CoV-2 ORF8 with SARS-CoV and SARS-CoV-2 ORF7a (PDB: 1xak and 6w37). Secondary structure assignments (blue cartoon arrows) correspond to the structures of SARS-CoV-2 ORF8 (top) and SARS-CoV-2 ORF7a (bottom). Cysteine residues involved in disulfide formation are highlighted (salmon). The conserved cysteine-cysteine linkages between ORF7a and ORF8 are shown (bottom:black) as well as the unique ORF8 cysteine-cysteine linkage (top:red). (B) Alignment of SARS-CoV-2 ORF8 and SARS-CoV ORF7a (PDB: 1xak). Alignment produced a DALI server Z-score=4.8 and an RMSD=2.4. Unique region of ORF8 structure is highlighted (light green). (C) ORF8 and ORF7a intramolecular disulfide bonds. The disulfides structurally conserved between the two proteins are shown as well as the ORF8-specific disulfide bond. (D) Alignment of CoV-2 ORF8 and other representative Ig-like fold proteins (PDBs: Dscam1, 4×83; CD244, 2ptu). Alignments produced a DALI server Z-score=6.2 & 6.1 and an RMSD=2.4 & 3.5, respectively.

SARS-CoV-2 ORF8 separates into two distinct species when analyzed by size exclusion chromatography. Comparison of the elution peak volumes with molecular weight standards suggest dimeric and monomeric forms (Fig. 3A). The asymmetric unit within the crystal contains the dimeric form (Fig. 3B) and exhibits imperfect two-fold non-crystallographic symmetry. The dimer is linked by an intermolecular disulfide bridge formed between two copies of Cys20 (Fig. 3B). Generation of the surface electrostatic potential for each monomer shows the interfaces are complementary (Fig. 3B). The dimeric interface amounts to approximately 1320 Å^2^ in buried surface area (*18*). The intermolecular bonds are primarily contributed by β1 and β7 residues and the loop joining β3 and β4 (Fig. 3C-D). The intermolecular disulfide bridge orchestrates the non-crystallographic symmetry axis (Fig. 3C). Val117 forms a hydrophobic interaction with its symmetry-related counterpart. This central hydrophobic region is flanked by salt bridges between Arg115 and Asp119 (Fig. 3C). The ends of the interface are stabilized by a “clamp” loop which makes main-chain hydrogen bonding interactions between Phe120 of one subunit and Ala51 and Arg52 of the other (Fig. 3D). The guanidino group of Arg52 in turn reaches across and forms a hydrogen bond with the carbonyl group of the Ile121 main-chain. An additional hydrogen-bond is formed between ε-amino group of Lys53 and Ser24. These features are near-identical on the opposite side of the interface, owing to the symmetry of the dimer.

**Fig 3.**
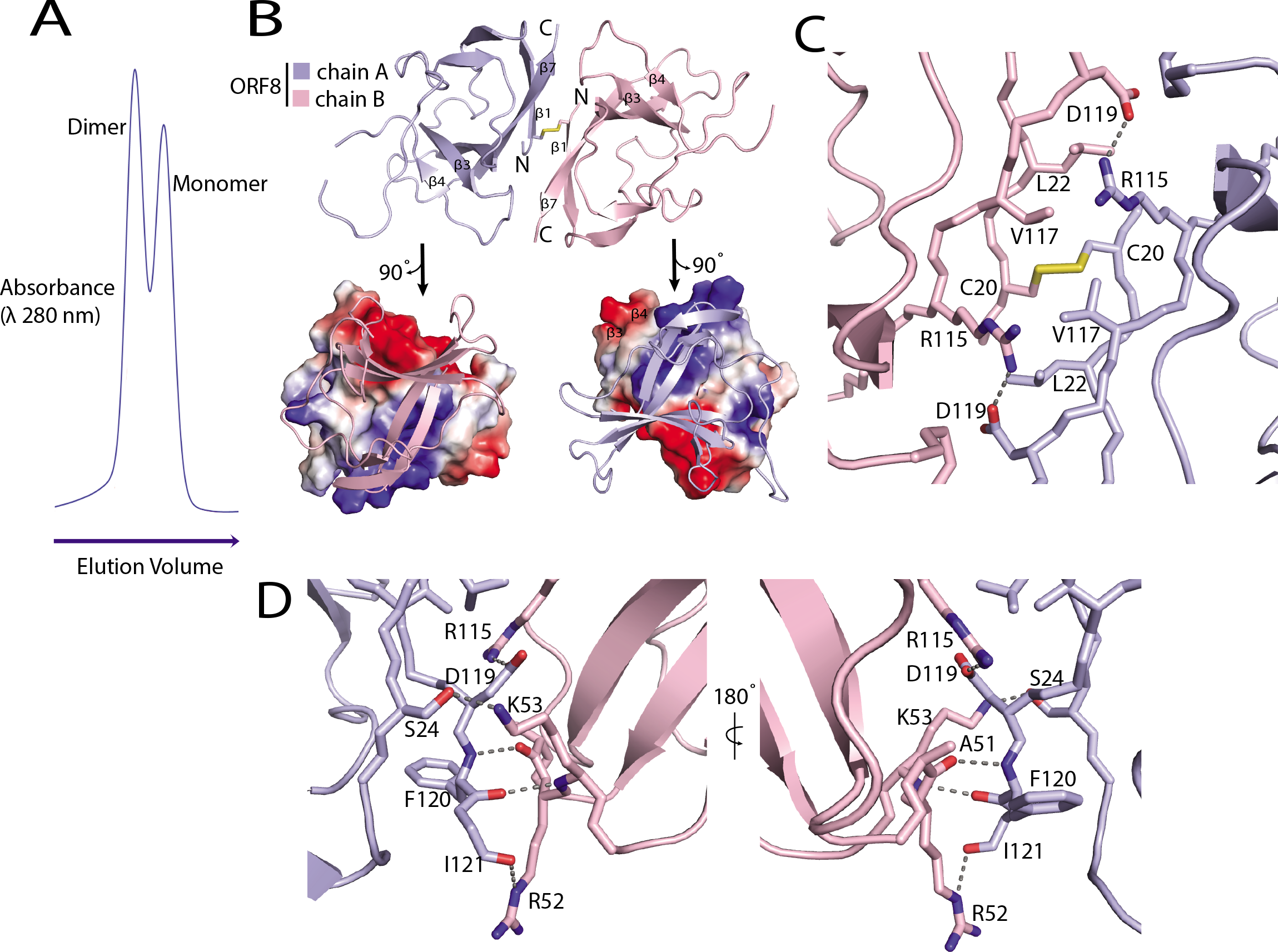
SARS-CoV-2 ORF8 forms a disulfide-linked homodimer. (A) Size exclusion chromatography (SEC) elution profile showing peaks corresponding to dimeric and monomeric ORF8 species. Absorbance is measured as a product of elution volume. (B) The top half of the panel communicates a cartoon representation of the asymmetric unit containing a single copy of the ORF8 dimer. The intermolecular disulfide bridge is formed between two cysteines, both corresponding to position 20 in the primary sequence. N- and C-termini are labeled accordingly. The bottom half shows an electrostatic potential surface representation of each monomer generated at neutral pH with positive, negative and neutral charges colored blue, red and grey respectively. (**C**) Detailed view of the dimeric interface, centered on the intermolecular disulfide bridge. (**D**) The edge of the dimeric interface is stabilized by multiple hydrogen-bonds. The opposite side of the interface displays a near-identical arrangement. Key residues are labeled, hydrogen bonds and salt bridges are shown as dashed lines.

Sequence alignment of SARS-CoV-2 ORF8 and its closely related bat betacoronavirus RaTG13 and SL-CoVZC45 orthologs revealed the six Cys forming intramolecular disulfide bridges are all conserved. These Cys are also conserved in SARS-CoV and most of its relatives (Fig. 4), and are also present in the corresponding regions of SARS-CoV ORF8a and b. However, Cys20, is not conserved in SARS-CoV ORF8 and bat viruses clustering phylogenetically with human SARS-CoV (Fig. 4). The residues immediately surrounding Cys20 are also conserved in the most recent bat precursors of SARS-CoV-2. The features thus required for the overall fold of the ORF8 monomer are well-preserved across SARS-CoV, SARS-CoV-2, and related betacoronavirus. The covalent dimer, is, however, an evolutionarily recent addition among human betacoronaviruses unique to SARS-CoV-2.

**Fig 4.**
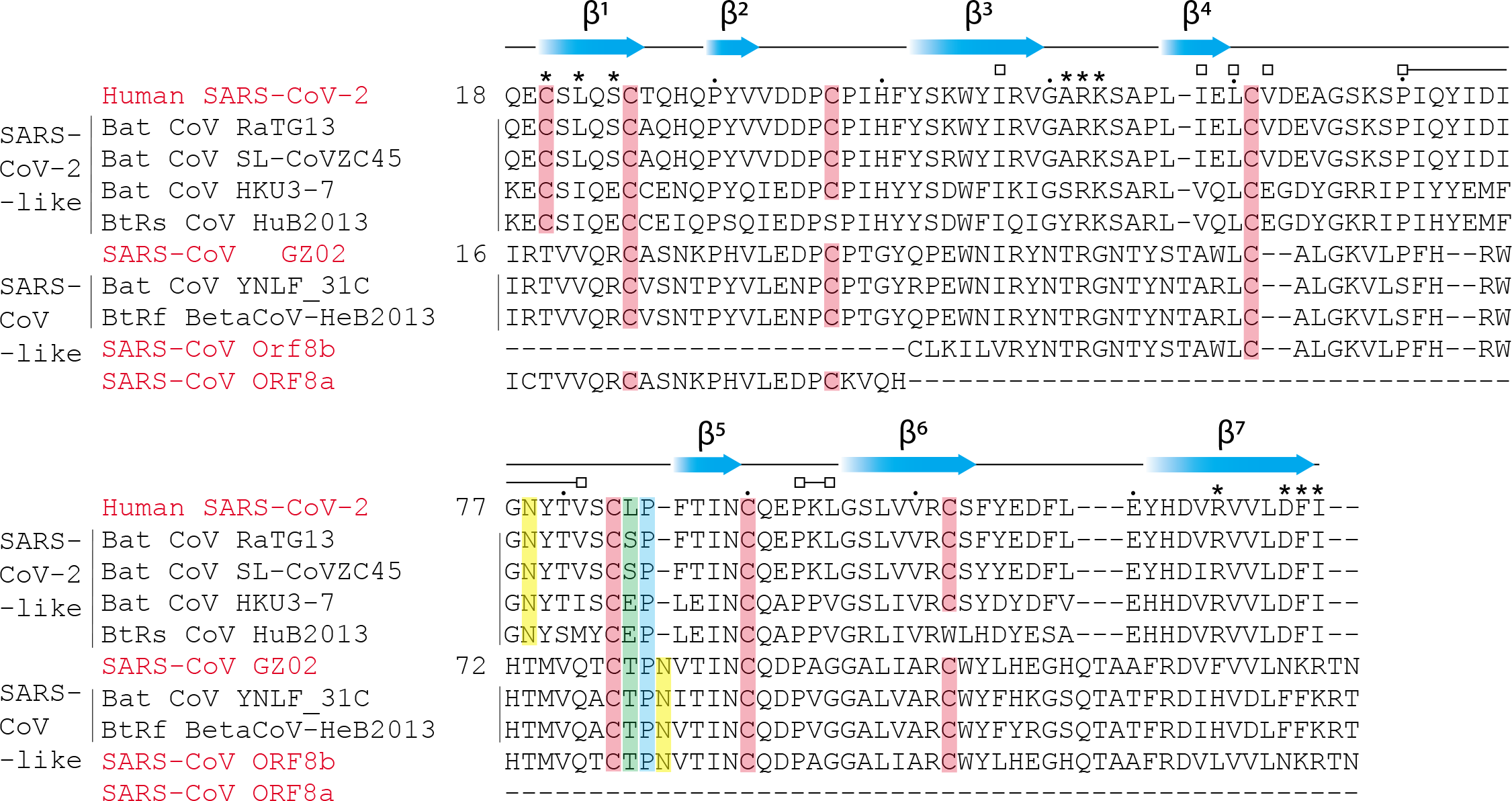
Conserved and unique features of SARS-CoV and SARS-CoV-2 ORF8. Primary sequence alignment of SARS-CoV-2 ORF8, SARS-CoV ORF8 and closely related ORF8 homologs found in bat betacoronavirus strains (*3*). Secondary structure assignments (blue cartoon arrows) correspond to the SARS-CoV-2 ORF8 structure. Putative and conserved biochemical features are highlighted: cysteines-disulfides (salmon), cis-proline (blue), residue 84 (green; SARS-CoV-2 ORF8 numbering), N-glycosylation site (yellow; predicted site for SARS-CoV-2) (*12*). Asterisks designate residues contributing to the ‘covalent’ dimer interface. Open squares designate residues contributing to the alternate dimeric interface. Dots designate intervals of ten amino acids according to SARS-CoV-2 ORF8 numbering.

Phylogenetic analysis of SARS-CoV-2 strains revealed two predominant isoforms of ORF8 in circulation, containing either Leu84 and Ser84 (*4, 7*). The structure reported here is of the Leu84 form. In the ORF8 structure, residue Leu84 is flanked by the disulfide-forming Cys83 on one side and by Pro85 on the other (Fig. 1C). Pro85 adopts the unusual *cis*-conformation. Both Cys83 and Pro85 are conserved amongst ORF8 orthologs. Given the position of Leu84, with its solvent exposed side-chain, it seems unlikely mutation would influence overall tertiary structure. Leu84 is also distal to both novel SARS-CoV-2-specific dimer interfaces. The biological role of residue 84 remains to be determined, but its unusual positioning controlled by a disulfide and cis-Pro together suggests its likely role in function.

Another major region of sequence unique to SARS-CoV-2 and its closest relatives begins immediately after Cys 61 and extends until just before the Cys83-Leu84-Pro85 conserved motif. A SARS-CoV-2-specific _73_YIDI_76_, motif occurs at the center of this unique region. The YIDI motif is responsible for stabilizing an extensive non-covalent dimer interface in the crystal, scored as highly significant (*18*) on the basis of its 1,700 Å^2^ of buried surface area and hydrophobicity. This suggests the non-covalent dimer seen in the crystal is a special feature of SARS-CoV-2 absent in SARS-CoV. The combination of Leu95, Ile58, Val49, and Pro56 form a hydrophobic interaction with Tyr73 of the YIDI motif (Fig 5C, 5D). Crystallographic contacts revealed an extensive array of hydrophobic interactions between residues 71-75 of chain A that interdigitate with the corresponding residues of a symmetry-related copy of chain B, distinct from the B-subunit of the covalent dimer (Fig 5E, 5F). The center of the interface comprises a two-stranded parallel β-sheet that is distinct from the core β-sheets of the monomeric Ig fold. Taken in combination with the Cys20-mediated covalent dimer interface, the structure shows two sequence regions unique to SARS-CoV-2 control the oligomerization and crystal packing of ORF8, and potentially mediate higher-order macromolecular assemblies unique to SARS-CoV-2 (Fig. 5G). These observations show how recent evolutionary changes in the sequence of SARS-CoV-2 relative to its more benign precursors could contribute to a unique higher-order assembly mediating unique functions in immune evasion and suppression.

**Fig 5.**
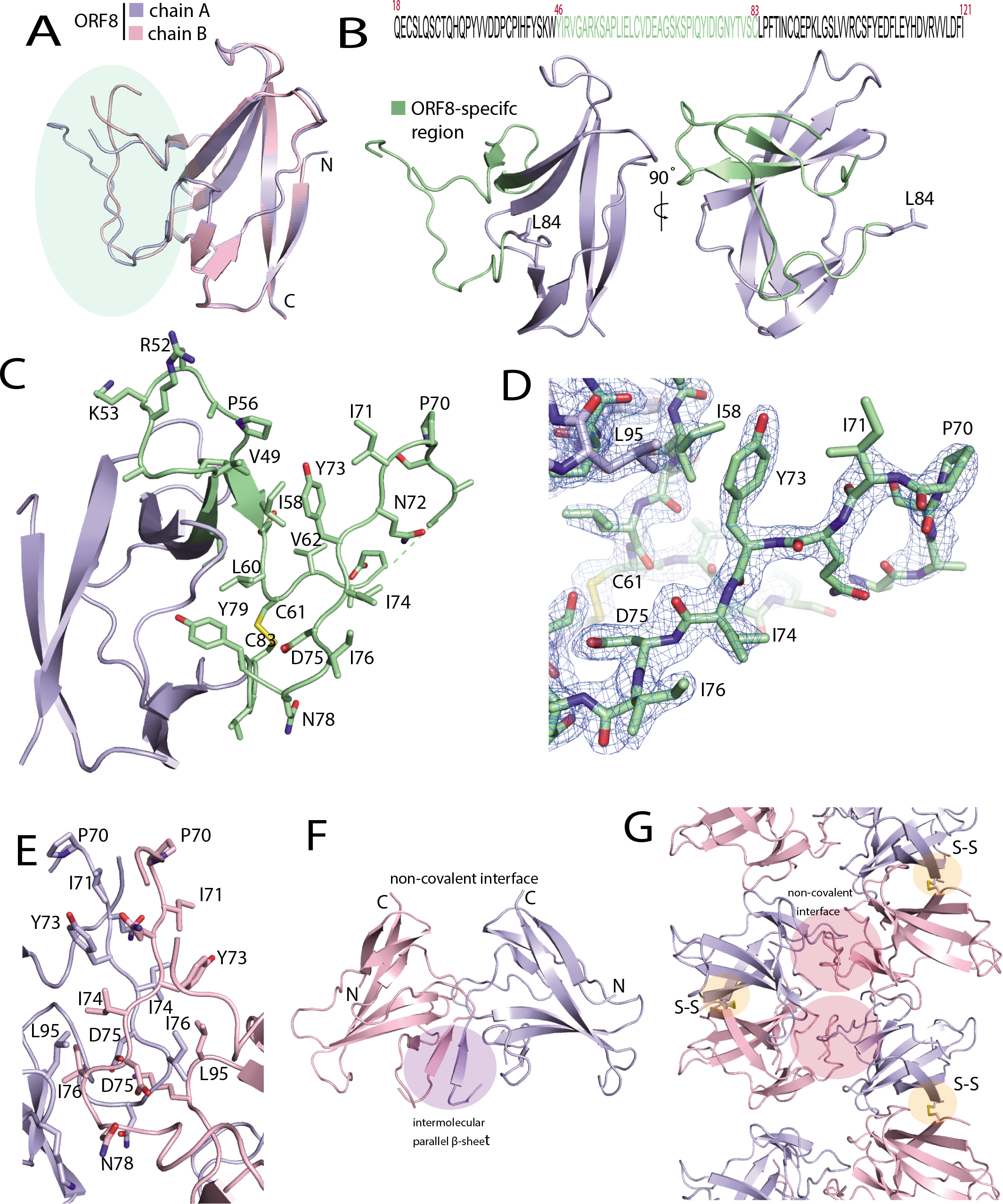
SARS-CoV-2 ORF8 contains a large, unstructured insertion. (A) Structural alignment of ORF8 chain A and B of the disulfide-linked dimer. The region corresponding to the ORF8-specific region is highlighted in green. (B) Primary sequence of the SARS-CoV-2 ORF8 construct used in this study is shown top panel. The ORF8-specific region is highlighted in green. Below is a cartoon representation of the monomer with the ORF8-specific region colored green. (C) A closeup of the ORF8-specific region is annotated. Notable residues are shown as sticks and labeled accordingly. (D) Stick representation of the insertion with 2Fo-Fc electron density map. The map is contoured at 2σ and represented as a blue mesh. (E) The crystallographic contact between ORF8 chain A and B form a non-covalent interface highlighted by an extensive array of hydrophobic residues. The residues are annotated and shown in stick form. (F) The non-covalent interface between ORF8 chain A and B forms a short, parallel β-sheet (G) Cartoon representation of alternating covalent disulfide and non-covalent interfaces in the ORF8 crystal lattice.

## Materials and Methods

### Protein expression and purification

The gene for wild-type SARS-CoV-2 ORF8 18-121 was subcloned from a cDNA kindly provided by D. Gordon and N. Krogan (UCSF) into the pET His6 TEV LIC cloning vector (2B-T). The plasmid was transformed into *E. coli* strain BL21(DE3) Rosetta pLysS (QB3 MacroLab, UC Berkeley), then expressed overnight (0.5 mM IPTG) at 20 °C in Luria Broth (LB) containing ampicillin and chloramphenicol. Cells were pelleted and resuspended in lysis buffer (50 mM Tris pH 8.0, 2 mM EDTA, 100 mM NaCl, 1 mM DTT, 0.5% Triton-X100) supplemented with protease inhibitors (Roche) and lysed by sonication. Lysate was clarified by centrifugation and the pellet washed with lysis buffer and sonicated for an additional 10 minutes to homogenize. Suspension was again clarified by centrifugation, the pellet resuspended in solubilization buffer (100 mM Tris pH 8.5, 6 M guanidine hydrochloride, 10 mM reduced glutathione) followed by Dounce homogenization and incubated at room temperature with rocking for 1 hour. Insoluble particulates were removed by centrifugation and the supernatant applied to nickel-charged agarose (GE Healthcare) preequilibrated in solubilization buffer. The resin was washed with solubilization buffer and bound His-tagged ORF8 eluted with elution buffer (100 mM Tris pH 8.5, 6 M guanidine hydrochloride, 10 mM reduced glutathione, 350 mM imidazole). Solubilized ORF8 was added drop-wise to a 50 fold excess of cold refolding buffer (50 mM Tris pH 8.0, 500 mM L-arginine, 2 mM EDTA, 5 mM reduced glutathione, 0.5 mM oxidized glutathione, 0.2 mM PMSF) over a period of 2 hours with gentle stirring followed by overnight incubation at 4 °C. The refolding solution was filtered, concentrated and applied to a Superdex S75 size-exclusion column equilibrated in buffer (20 mM Tris-HCl, pH 8.0, 150 mM NaCl, 1 mM EDTA). Folded ORF8 eluted as two peaks corresponding to monomer and dimer. Both peaks were pooled and incubated with TEV protease overnight. Cleavage products were passed through nickel-charged agarose (GE Healthcare) to remove any uncleaved ORF8. The flow-through was concentrated and applied to a Superdex S75 size-exclusion column equilibrated in buffer (20 mM Tris-HCl, pH 8.0, 150 mM NaCl, 1 mM EDTA). Both monomer and dimer peaks were pooled and concentrated.

The plasmid containing the ORF8 V32M, L84M double mutant coding sequence was generated by QuikChange site-directed mutagenesis according to the manufacturer’s instructions (Agilent). Selenomethionine labeled ORF8 double mutant protein was expressed using SelenoMet Medium Base according to manufacturer’s instructions (Molecular dimensions). Purification was carried out in the same manner as wild-type but the TEV-cleavage step was omitted.

The expression construct has been made available at addgene.org.

### X-ray crystallography

Crystals of wild-type ORF8 were grown using the hanging-drop vapor-diffusion method at 18 °C. 2 μl of the protein sample (7.8 mg ml-1) was mixed with 2 μl of reservoir solution and suspended over a 500-μl reservoir of 100 mM sodium dihydrogen phosphate pH 6.5, 12 % (w/v) PEG8000. Crystals appeared after 5 days and continued to grow for approximately one week. Crystals were transferred into cryoprotectant (100 mM sodium dihydrogen phosphate pH 6.5, 12 % (w/v) PEG8000, 20 mM Tris pH 8.0, 150 mM NaCl, 1 mM EDTA, 30 % (v/v) glycerol) and flash-frozen by plunging into liquid nitrogen. A native dataset was collected from a single crystal of space group *P*4_1_2_1_2 under cryogenic conditions (100 K) at a wavelength of 1.00001 Å using a Dectris Pilatus3 S 6M detector (Beamline 5.0.2, Advanced Light Source, Lawrence Berkeley National Laboratory).

Crystals of selenomethionine (SeMet)-labeled V32M/L84M ORF8 were produced using the same reservoir condition as wild-type and a lower concentration of protein (2.5 mg ml-1). Crystals exhibiting a different morphology to wild-type appeared after 2 days and continued to grow for approximately 4 days. A single-wavelength anomalous dispersion (SAD) selenium peak dataset in space group *I*4_1_22 was collected at a wavelength of 0.97903 Å.

The diffraction data were indexed and integrated using XDS (*19*). Integrated reflections were scaled, merged and truncated using the CCP4 software suit (*20*). Initial phases were obtained for the SeMet dataset using the Phenix autosol pipeline (*21*). This solution led to a map that enabled the full ORF8 main-chain to be traced which was partially refined and in then used as a search model for molecular replacement with respect to the native dataset using PHASER (*22*). The space group of the native dataset is *P*4_1_2_1_2 with one ORF8 dimer in the asymmetric unit. Iterative rounds of manual model building and refinement were carried out using Coot (*23*) and Phenix Refine (*21*) respectively. Statistics of the final native structure are shown in Table 1. Structural figures were produced using the program PyMOL (https://pymol.org).

**Table 1.** Data collection and refinement statistics

|  | SARS-CoV-2 ORF8<br>(PDB code: 7JTL) |  |
| --- | --- | --- |
| Data collection |  |  |
|  | Native | SeMet |
| Space group | *P*4<sub>1</sub>2<sub>1</sub>2 | *I*4<sub>1</sub>22 |
| Cell dimensions |  |  |
| *a*, *b*, *c* (Å) | 44.258 44.258 264.111 | 49.21 49.21 270.65 |
| $\alpha$, $\beta$, $\gamma$ (°) | 90, 90, 90 | 90, 90, 90 |
| Resolution (Å) | 43.65-2.04 (2.113-2.04) | 48.42-2.92 (3.10-2.92) |
| *R*<sub>pim</sub> | 0.0323 (0.5549) | 0.020 (0.498) |
| *I* / σ*I* | 14.92 (1.20) | 27.9 (2.8) |
| Completeness (%) | 97.8 (90.2) | 99.9 (99.8) |
| Redundancy | 10.0 (7.6) | 27.4 (28.8) |
| Refinement |  |  |
| Resolution (Å) | 43.65-2.04 |  |
| No. reflections | 17005 (1399) |  |
| *R*<sub>work</sub> / *R*<sub>free</sub> | 0.2199 (0.3516)/ 0.2640 (0.3844) |  |
| No. atoms |  |  |
| Protein | 1609 |  |
| Ligand/ion | 0 |  |
| Water | 201 |  |
| *B*-factors |  |  |
| Protein | 45.36 |  |
| Ligand/ion |  |  |
| Water | 44.28 |  |
| R.m.s. deviations |  |  |
| Bond lengths (Å) | 0.008 |  |
| Bond angles (°) | 1.00 |  |

Values in parentheses are for highest-resolution shell. Redundancy for SeMet data is shown for I<sup>+</sup> and I<sup>-</sup> separately

### Sequence alignment

Primary sequences of ORF8 and ORF7 from human and bat isolates were collected from GenBank. Sequences used for alignment were selected based on previous phylogenetic analyses (*12*). Primary sequence alignment of ORF8 homologs was performed using the ClustalOmega server (*24*). Sequence alignment of ORF7a and ORF8 and secondary structure annotations were guided by DSSP through the DALI server (*25*) and manually adjusted based on visual inspection of the structures.

## Acknowledgements

This work was supported by UCOP emergency seed grant R00RG2347 (JHH), and National Institutes of Health grants R37 AI112442 (JHH), R01 AI120691 (XR and JHH), P50 AI150476 (JHH), F32 AI150495 (CZB), and F32 AI152971 (RMH). The Berkeley Center for Structural Biology is supported in part by the Howard Hughes Medical Institute. The Advanced Light Source is a Department of Energy Office of Science User Facility under Contract No. DE-AC02-05CH11231. The ALS-ENABLE beamlines are supported in part by the National Institutes of Health, National Institute of General Medical Sciences, grant P30 GM124169.

## Coordinates

Coordinates have been deposited in the PDB and are available as entry 7JTL.

## Notes

### Competing Interest Statement

The authors have declared no competing interest.

